# Modulation of prefrontal couplings by prior belief-related responses in ventromedial prefrontal cortex

**DOI:** 10.1101/2023.07.25.549989

**Authors:** Bin A. Wang, Sabrina Drammis, Ali Hummos, Michael M. Halassa, Burkhard Pleger

## Abstract

Humans and animals can maintain constant payoffs in an uncertain environment by steadily re-evaluating and flexibly adjusting current strategy, which largely depends on the interactions between the prefrontal cortex (PFC) and mediodorsal thalamus (MD). While the ventromedial PFC (vmPFC) represents the level of uncertainty (i.e., prior belief about external states), it remains unclear how the brain recruits the PFC-MD network to re-evaluate decision strategy based on the uncertainty. Here, we leverage nonlinear dynamic causal modeling on fMRI data to test how prior belief-dependent activity in vmPFC gates the information flow in the PFC-MD network when individuals switch their decision strategy. We show that the prior belief-related responses in vmPFC had a modulatory influence on the connections from dorsolateral PFC (dlPFC) to both, lateral orbitofrontal (lOFC) and MD. Bayesian parameter averaging revealed that only the connection from the dlPFC to lOFC surpassed the significant threshold, which indicates that the weaker the prior belief, the less was the inhibitory influence of the vmPFC on the strength of effective connections from dlPFC to lOFC. These findings suggest that the vmPFC acts as a gatekeeper for the recruitment of processing resources to re-evaluate the decision strategy in situations of high uncertainty.

**Author Summary:** Prefrontal cortex (PFC) together with the mediodorsal thalamus (MD) jointly establish computations critical for behavioral adaptations. While the task uncertainty (i.e., prior belief) was represented by the ventromedial PFC (vmPFC), it remains unclear how the PFC-MD network reallocates the processing resources to re-evaluate decision strategy under uncertainty. Here we filled this gap by leveraging the Bayesian hierarchical modelling and nonlinear dynamic causal modelling in an associative learning task. We found that in situations of high uncertainty, the prior belief-related responses in vmPFC significantly strengthened effective connectivity from the dorsolateral PFC to the orbitofrontal cortex, but not to the MD. The findings provide evidence for the role of vmPFC in driving the re-evaluation of the decision strategy during behavioral adaptations in situations of uncertainty.

## Introduction

The prefrontal cortex (PFC) consists of several regions that are thought to play an important role in flexible decision-making. The dorsolateral PFC (dlPFC) is assumed to support executive functions (Jones & Graff-Radford, 2021), whereas the orbitofrontal cortex (OFC) appears to be involved in the flexible adaptation of behavior (Schoenbaum et al., 2021; Wang et al., 2023). The ventromedial PFC (vmPFC) was shown to be associated with the estimation of the value and saliency of sensory events and thereby guides value-based decision-making (Dundon et al., 2021). Interactions between these regions are thought to implement a variety of functions relevant to the flexibility by which cognitive resources are deployed. Interaction between medial PFC and the anterior cingulate cortex, for instance, are thought to contribute to the updating of beliefs about higher-order contextual associations (M. Botvinick et al., 1999; M. M. Botvinick et al., 2001; Sarafyazd & Jazayeri, 2019), whereas the interplay between OFC and vmPFC appears important for the prediction of value-based behavioral changes (Howard et al., 2016).

Recent studies in animals (Halassa & Kastner, 2017; Mukherjee et al., 2021; Saal et al., 2017; Schmitt et al., 2017) and humans (Hwang et al., 2017; Wen et al., 2021) revealed compelling evidence that cognitive flexibility also depends on interactions between distinct PFC subregions and the mediodorsal nucleus of the thalamus (MD). These studies have provided a complementary perspective on thalamic function challenging the classical notion of the thalamus as a sensory relay (Sherman, 2016). Combining hierarchical Bayesian modeling with fMRI in humans, we recently unveiled distinct prefrontal connections targeting the MD in relation to the participant’s prior belief during associative learning (Wang & Pleger, 2020). The surprise about an unexpected outcome lowered the prior belief about the cue-target association and hence triggered a switch of the decision strategy through modulations of connections between MD and lateral OFC (Wang & Pleger, 2020). These findings are supported by neuronal recordings obtained from mice. When mice decided between different sets of learned cues that directed attention to either visual or auditory targets, responses from the medial PFC reflected both, the individual cue as well as its importance as a task-rule (Rikhye et al., 2018). The MD, on the other hand, appeared to facilitate switching between cueing contexts by supporting or suppressing task-associated representations in the PFC (Mukherjee et al., 2021; Rikhye et al., 2018; Schmitt et al., 2017). Together, these findings from mice and humans emphasize crucial prefrontal-MD computations necessary for learning stimulus-incentive associations.

We recently refined a previously developed computational prefrontal-MD model, inspired by cell recordings obtained from mice (Rikhye et al., 2018), and trained it on human empirical data to test whether the re-evaluation and adjustment of the decision strategy in both species follow the same computational principles (Hummos et al., 2022). We found that the MD learned abstract representations of its cortical inputs through biologically plausible Hebbian learning rules. Direct feedback from MD to prefrontal cortex supported switching between behavioral strategies, while lateral OFC (lOFC) constantly accumulated evidence for a strategy switch based on rapid Bayesian estimation. Following these computational rules, our human fMRI results revealed that lOFC directed its outputs to MD, rendering MD as the brain site which dynamically integrates crucial inputs relevant for forming the behavioral strategy. These abstract MD representations and their ability to reorganize prefrontal computations describes an efficient way how the brain utilizes the MD to integrate inputs from other brain regions and to dynamically select between competing behavioral strategies (Hummos et al., 2022).

Another prefrontal region, the vmPFC, was shown to encode the beliefs about outcome values, which represents an intermediate signal required for efficient prefrontal-MD computations underpinning the re-evaluation of decision strategy. In our prefrontal-MD model (Hummos et al., 2022), we directed the vmPFC output on the executive dlPFC, as the most relevant receiver of value-related information, but this dlPFC-vmPFC interaction was not directly supported by empirical evidence. Using bilinear Dynamic Causal Modelling (DCM), we could not directly model the vmPFC as an additional hub in the prefrontal-MD network since the vmPFC was not among the regions involved in decision switches. In the present study, we therefore applied non-linear DCM, which allowed us to capture the non-linear history of prior synaptic activity (Stephan et al., 2008), and hence the modulatory (second-order) effects of prior belief-dependent responses in vmPFC on the gain of activity within the prefrontal-MD network. We re-analyzed our previously collected fMRI dataset and combined Bayesian hierarchical modeling with nonlinear DCM on fMRI data to test how prior belief-related activity in the vmPFC tunes functional couplings in the dlPFC-lOFC-MD network during the adjustment of the decision strategy.

## Results

The participants’ behavioral performance during the associative learning task has been presented in more detail in our previous studies (Hummos et al., 2022; Wang et al., 2020; Wang & Pleger, 2020). In brief, we found a significant effect of learning (one-way ANOVA: F (4,27) =256, p < 0.001) and post-hoc paired t-tests revealed that participants made significantly more correct predictions in learning blocks with high predictability (i.e., 90%/10%) than in blocks with low predictability (i.e., 70%/30%, t(1,27) = 20.75, P < 0.001, Bonferroni-corrected) or unpredictability (i.e., 50%, t(1,27) = 24.70, p < 0.001, Bonferroni-corrected). In addition, we found that the participants had a lower prior belief and required more time to respond (p < 0.001) in trials when they changed their decision strategy (*Switching*), compared to the trials where the decision strategy did not change (*Staying*).

To assess the parametric modulation by the prior beliefs, both 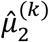 and 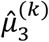 derived from HGF model were included as modulatory parameters in the GLM of fMRI data. We found that responses from the vmPFC reflected the prior belief about the sample-target contingency (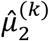, Fig. 1C, x = -6, y = 62, z = -4, t(31) = 6.69, p < 0.05, FWE whole-brain corrected). Besides the vmPFC, also the left precuneus (x = -10, y = -56, z = 26) and middle cingulate cortex (x = 2, y = -12, z = 38) represented significant prior belief-related activity (p < 0.05, FWE whole-brain corrected). The analysis of parametric effects related to prior belief about volatility 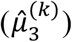 did not reveal any significant brain regions (p > 0.05, FWE whole-brain corrected).

**Fig. 1.**
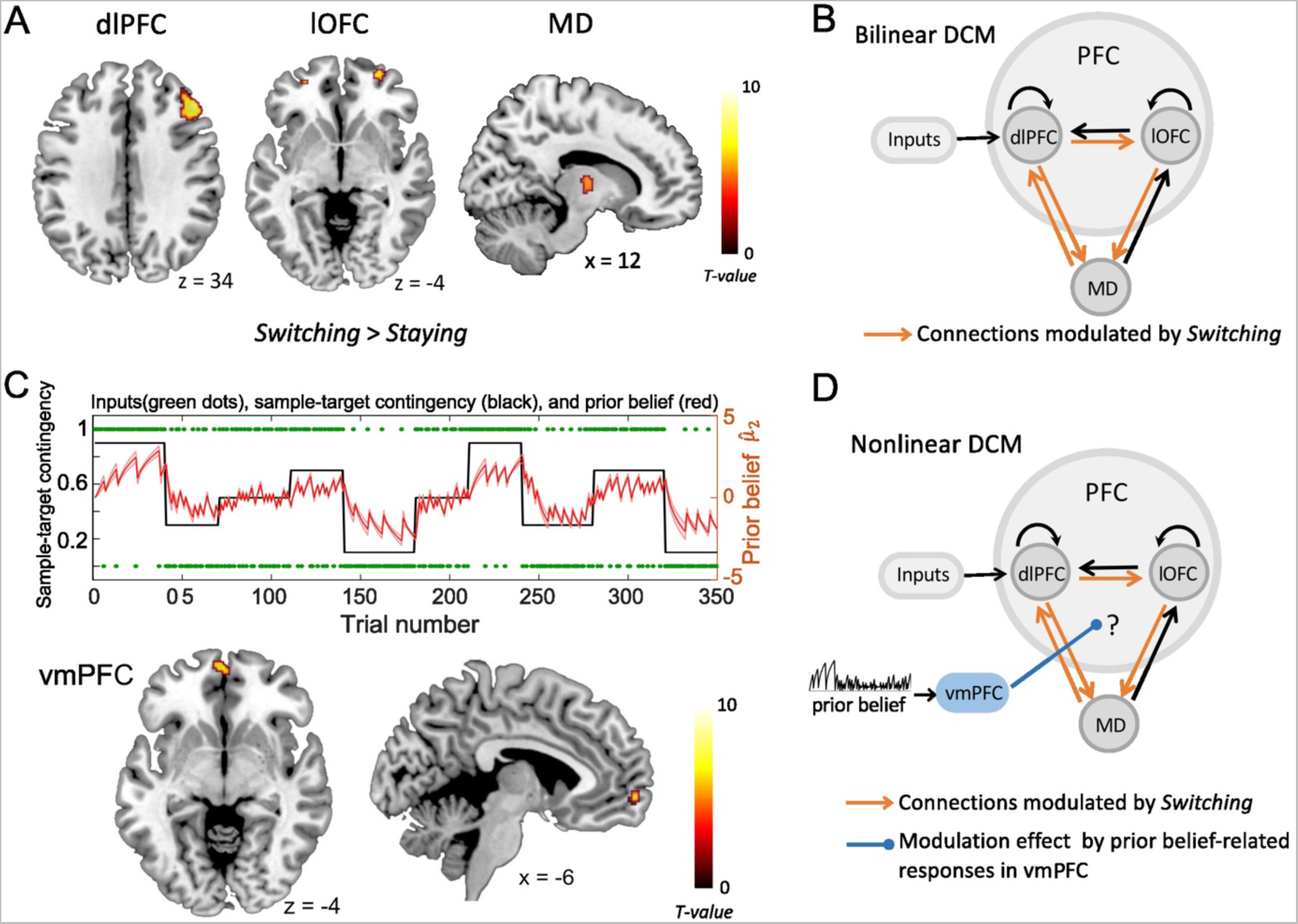
The fMRI activity and dynamic causal modeling (DCM). **A**. Strategy switches (*Switching>Staying*) entailed significant BOLD activity in right dlPFC, right lOFC and right MD (Hummos et al., 2022). **B**. The bilinear DCM revealed that four connections (orange), i.e., from lOFC to MD, from dlPFC to lOFC, as well as between dlPFC and MD in both directions were significantly strengthened by strategy switches (*Switching* > *Staying*) (Hummos et al., 2022). The black connections indicate endogenous connections between brain areas. **C**. Upper panel: The prior beliefs over the course of the experiment. The red shaded area indicates the standard error of the mean (SEM) of prior beliefs over time. Lower panel: Prior belief-related vmPFC activity is projected on sagittal and axial MRI brain slices (p < 0.05, whole brain-FWE corrected). **D**. The non-linear DCM, which we used to test how prior belief-related activity in the vmPFC (blue) gated the information flow in the dlPFC-lOFC-MD network. (dlPFC–dorsolateral prefrontal cortex; lOFC–lateral orbitofrontal cortex; MD–thalamic mediodorsal nucleus; vmPFC-ventromedial prefrontal cortex)

As shown in our recent study (Hummos et al., 2022), the Bayesian model comparison across different plausible bilinear dlPFC-lOFC-MD models revealed that the adjustment of the decision strategy (*Switching*) significantly modulated connections from lOFC to MD, from dlPFC to lOFC, as well as between dlPFC and MD in both directions (Fig. 1B). In the present study, using nonlinear DCM, we tested how prior belief related response from vmPFC modulated the connection strength in the dlPFC-lOFC-MD network (Fig. 1D). More specifically, we tested which two of the four projections were directly modulated by vmPFC activity. This resulted in six competing models that were further evaluated with Bayesian model selection (BMS) (Supplementary Fig. S1A). BMS revealed that the prior belief associated activity in vmPFC had a modulatory influence on the projections originating in the dlPFC and targeting both, lOFC and MD (Supplementary Fig. S1B and Fig. 2A). The posterior probability for the winning model was 0.48, surpassing the posterior probabilities of the other tested models which ranged from 0.01 to 0.25 (Supplementary Fig. S1B). This winning nonlinear model assumed a direct effect of prior belief on vmPFC whose activity then mediated the gain of the dlPFC→lOFC and dlPFC→MD connections. We next applied Bayesian parameter averaging (BPA), which computes a joint posterior density for the entire sample and found that the prior belief-dependent vmPFC activity significantly influenced the dlPFC→lOFC connection (posterior probability = 0.94, Fig. 2B). The modulations of the dlPFC→MD connection showed a trend into the same direction, but the posterior probabilities failed to surpass the 90% threshold (posterior probability = 0.81, Fig. 2B). The modulatory effect was generally inhibitory, suggesting that the weaker the activity in vmPFC, and the lower the prior belief (i.e., high task uncertainty), the stronger was the connection strength from dlPFC to lOFC.

**Fig. 2.**
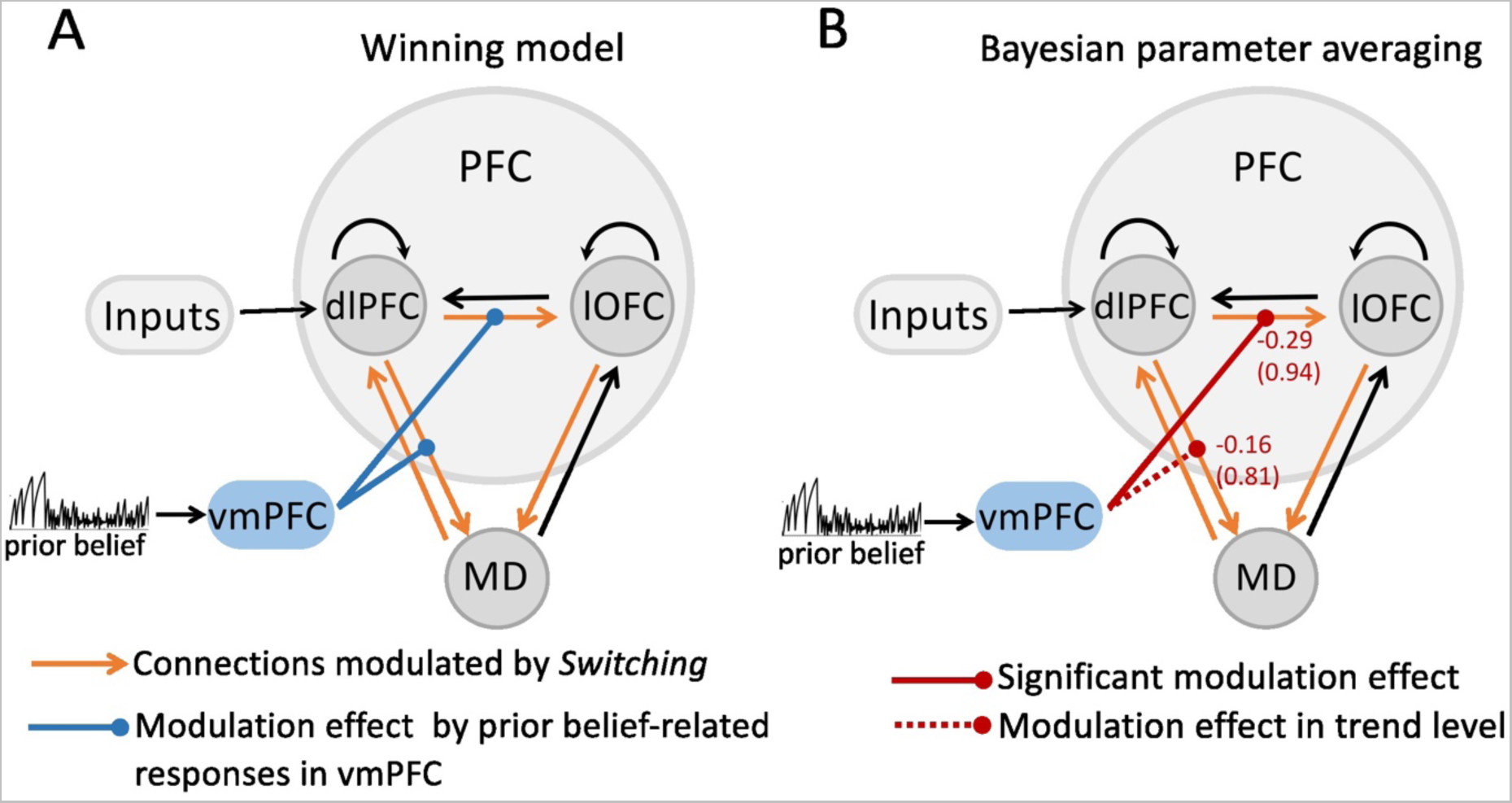
The modulatory effect on the dlPFC-lOFC-MD network by prior belief-associated vmPFC activity. **A**. The Bayesian Model Selection across the possible six models revealed the winning model in which prior belief-dependent vmPFC activity specifically influenced the connections originating in the dlPFC and targeting both, lOFC and MD. The black connections indicate endogenous connections between brain areas. **B**. Bayesian parameter averaging revealed that the prior belief-dependent vmPFC activity significantly inhibited the dlPFC→lOFC connection (posterior probability = 0.94, solid red line) suggesting that the higher the activity in vmPFC, and the stronger prior belief, the weaker was the connection strength from dlPFC to lOFC. The negative numbers shown next to the projections and above the posterior probabilities index the inhibitory inputs per second (Hz). The dlPFC→MD connection showed a trend into the same direction, but posterior probabilities failed to surpass the 90% threshold (posterior probability = 0.81, dotted red line).

## Discussion

In this study, we investigated how prior belief-related activity in the vmPFC gates the information flow in the dlPFC-lOFC-MD network underpinning the re-evaluation of decision strategy during associative learning. We showed that the strength of connections from dlPFC to lOFC fluctuated from trial to trial in relation to the level of task uncertainty, i.e., the prior beliefs about the cue-target contingency, signaled by inhibitory inputs from the vmPFC. The weaker the activity in vmPFC, and the stronger the task uncertainty (i.e., low prior belief), the stronger was the strength of effective connection from executive dlPFC to lOFC. These findings suggest that in situations with high task uncertainty, there is an increase in lOFC responses to inputs from dlPFC, which provides direct empirical evidence for the key role of prior belief-dependent synaptic plasticity in driving the re-evaluation of the decision strategy during flexible decision-making.

Representations of sensory stimuli are modulated by internal states and beliefs about the world, which are crucial for flexibly adjusting perception and behavior in humans and animals (Akrami et al., 2018; Snyder et al., 2015). The prior beliefs have been shown to warp neural representations in the frontal cortex, which allows the mapping of sensory inputs to motor outputs and to incorporate prior statistics in accordance with Bayesian inference (Sohn et al., 2019). These results uncover a simple and general principle whereby prior beliefs exert their influence on behavior by sculpting cortical latent dynamics. The failure of beliefs about the environmental state forms the prediction error and updates expectations for upcoming stimuli and associated rewards. In a previous study, we investigated how humans apply probabilistic computations, following Bayesian rules, to infer on joint brain representations of prior belief and decision switches (Wang et al., 2020). We found, that during such switches prior belief specifically modulated connectivity among the anterior insular cortex (AIC), the premotor cortex (PMd), and the inferior parietal lobule (IPL).

On a trial-by-trial basis, prior belief weakened connectivity between AIC and IPL when the sensory stimulus was expected, whereas it strengthened connectivity between AIC and PMd when the stimulus was unexpected (Wang et al., 2020). AIC has been shown to act as a core hub modulating the interaction of bodily, attentional, and anticipatory sensory signals (Allen et al., 2016; Craig, 2009; Sridharan et al., 2008). Our results furnish a picture in which AIC, in conjunction with other brain regions, contributes not only to the coordination of expectation and sensory inputs, but also to the integration of priors and prediction outcomes for updating beliefs specifically supporting strategy switches during associative learning. The present findings extend the scope of belief-related brain functions by the vmPFC. According to our non-linear DCM, vmPFC plays a key role in exerting modulatory influences on prefrontal interactions, which does not directly reflect prior belief such as the aforementioned AIC-network (Allen et al., 2016), but which critically depends on prior belief-dependent information from the vmPFC to flexibly guide strategy switches.

It has been shown that vmPFC plays a major role in sensory integration to achieve abstract and conceptual interpretation of the environment (Petrides & Pandya, 2007). The vmPFC represents signals more suited for subject-centered, internally driven motivational processes, whereas lOFC encodes signals for evaluating environment-centered, externally driven motivational processes (Bouret & Richmond, 2010). Functional interactions between vmPFC and lOFC in humans are thought to implement important functions relevant to cognitive flexibility and the prediction of value-based behavioral changes (Howard et al., 2016). In primates, the dlPFC has been shown to be anatomically (Mackey & Petrides, 2010; Saleem et al., 2014) and functionally (Kahnt et al., 2012) connected to the lOFC, and also in rodents, mPFC terminals -i.e., homologue of the primate dlPFC -were detected in the lateral and ventrolateral OFC (Dalley et al., 2004). Rodent mPFC projection neurons furthermore target the thalamus MD to regulate adaptive control to flexibly optimize behavioral responses in goal-directed behavior and receive an MD innervation (Carlén, 2017; Hoover & Vertes, 2007). Together, these anatomical evidence from rodents and non-human primates provide a solid anatomical basis for our winning nonlinear DCM model, in which the responses of lOFC to the presynaptic input from dlPFC depend on the history of inputs that they receive from vmPFC. Inconsistencies in the nomenclature and anatomical boundaries of PFC areas have made it difficult to compare data and interpret findings across species, especially between primates and rodents. In addition, we cannot exclude that the modulatory effect on the PFC interactions is implemented through an intermediate region, for example the amygdala or the striatum, given the fact that both receive many cortical inputs from prefrontal areas (Chang & Grace, 2018; Middleton & Strick, 2002).

The weaker the prior belief related activity in vmPFC, the stronger is the connection strength of the dlPFC projections, suggesting that the re-evaluation of decision strategy under high task uncertainty requires further processing resources controlled by vmPFC. This gating mechanism agrees well with the theoretical accounts of the free-energy principle (Friston, 2009). Accordingly, perception optimizes predictions by minimizing free energy with respect to perceptual inference, memory, attention and salience (Friston, 2010). Based on these principles, the high prior belief about the environmental state in our study may have weakened the strength of the dlPFC-output projections to minimize the engagement of additional cognitive resources and hence to lower the energy costs for the decision process. The process that alters synaptic strengths with time constants in the range of milliseconds to minutes, i.e., the so-called “short-term synaptic plasticity” (STP), is proposed to be the underlying neurobiological mechanisms for this nonlinear modulatory effect, including NMDA-controlled rapid trafficking of AMPA receptors (Diering & Huganir, 2018), synaptic depression/facilitation (Zucker & Regehr, 2002) or “early long-term potentiation (LTP)” (Frey & Morris, 1998).

There are limitations that should be considered when interpreting the current findings. First, single unit recording in mice are well suited to assess how distinct neuronal populations in subregions of the PFC and MD nuclei respond to specific task-related cues and contexts (Rikhye et al., 2018). FMRI recordings instead represent comparably crude measures of neural activity capturing hemodynamic responses only from large neuronal populations. These topographic inaccuracies, especially for signals originating in small cortical and subcortical structures, represent a general limitation of fMRI. Second, we did not split the MD into its subnuclei when we constructed the network models. A recently developed MRI-based atlas provides masks for four of these subregions (the medial (MDm), central (MDc), dorsal (MDd), and lateral (MDl)), created with connectivity-based methods applied to the high-resolution data from the Human Connectome Project (Li et al., 2022). In the current study with lower functional MRI resolution and signal-to-noise ratio, it remained difficult to reliably capture these subregions. Therefore, we considered the MD as a single region of interest. Considering these topographic inaccuracies, the trend level effect we found for the modulatory influence on the dlPFC-MD connection should be interpreted with caution. Third, although we focused on thalamocortical connections, other regions of the broader PFC-MD network, including the basal ganglia or the amygdala, could have also been involved in guiding cognitive flexibility. For example, the cortical-basal ganglia-thalamocortical subnetworks have been shown to be involved in impaired cognitive flexibility in patients with obsessive-compulsive disorder (Kim et al., 2022). Given the above limitations, it will be one of the major challenges for future decision-making research to extend the here proposed PFC-MD network by other key regions in the PFC, basal ganglia, and other subcortical structures, through combining various imaging techniques and cross-species approaches.

## Materials and Methods

### Participants and associative learning task

The analyses in this study were based on the previously collected dataset from twenty-eight healthy human participants (mean age ± SD: 25.3 ± 3.9 years, only male participants). The study was approved by the local ethics committee of the Ruhr-University Bochum. All participants gave written informed consent prior to participation. Demographics and the experimental design were described in more detail elsewhere (Hummos et al., 2022; Wang et al., 2020; Wang & Pleger, 2020).

In each trial, participants first received one out of two tactile cues for 500ms to the tip of their right index finger using an MRI-compatible Braille piezo stimulator with 8-pins (2×4 array) (Metec, Stuttgart, Germany). Subsequently, they had to predict (within 1300ms) whether the following tactile stimulus (i.e., target) will show the same pattern (e.g., 4 upper pins lifted) or the alternative pattern (4 lower pins lifted) by pressing one of two buttons (LumiTouch keypads, Photon Control) with the index or middle finger of the left hand. After an interval of 500-1500ms (jitter), the target was presented for 500ms which indicated whether the preceding prediction was correct or incorrect. A variable delay of 1500-3000ms separated trials. The predictability of the target stimulus was modulated by the strength of the cue–target contingency over time (i.e., strongly predictive blocks (90% and 10%), moderately predictive (70% and 30%), and nonpredictive (50%) blocks; either 30 or 40 trials per block to avoid predictability of the block onsets). The order of blocks was pseudorandomized and fixed across participants to ensure inter-subject comparability of the learning process. The fMRI experiment consisted of 350 trials in total, which were split into three runs, each lasting about 10 min.

In order to examine the flexibility in decision-making, we tested the adjustment of the decision strategy, i.e., whether the statistical property of the environment (sample stimulus matches or mismatches target stimulus) has been changed or not. To this end, we separated the trials across all sample–target associations into two conditions: 1) *Switching* condition (122 ± 10 trials): trials with decision switches across two successive trials, and 2) Staying condition (211 ± 17 trials): trials without decision switches across two trials.

### Estimating the prior belief with the Hierarchical Gaussian Filter

Human behavioral data were applied to a three-level Hierarchical Gaussian Filter (HGF) model using the HGF toolbox (v5.2) as implemented in TAPAS (Translational Algorithms for Psychiatry-Advancing Science, https://www.tnu.ethz.ch/en/software/tapas, Frässle et al., 2021), to calculate the individual trial-wise prior belief about external states at different levels. The first level of the HGF represents a sequence of inputs about the environmental states (i.e., whether the sample stimulus matches the target stimulus or not), the second level represents the sample-target contingency (i.e., the conditional probability, in logit space, of the target stimulus given the sample cue), and the third level represents the log-volatility of the environment. Each of these hidden states is assumed to evolve as a Gaussian random walk, such that its variance depends on the state at the next higher level (Mathys et al., 2011).

In the HGF, at any level *i* of the hierarchy, the prior belief about the external state on trial 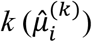 is evolved from the posterior belief of the previous trial (*k*-1)

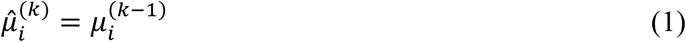

The posterior belief on trial 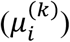 is updated based on the prediction error from the level below 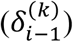 weighted by the precision of prediction 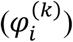:

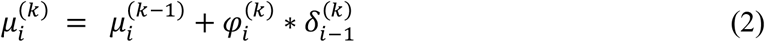

The precision of prediction 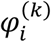 is updated with every trial and can be regarded as equivalent to a dynamic learning rate in reward learning models, as follows:

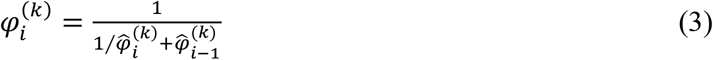

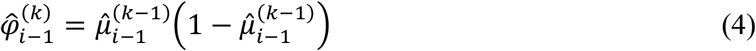

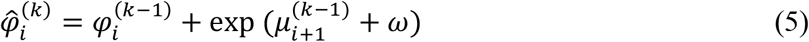

where *ω* is a free parameter of the perceptual model in HGF, which determines the step size between consecutive time steps. The prediction error 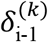, which drives learning at the second level of our HGF model, is defined as the difference between the actual outcome and its estimated probability before the outcome:

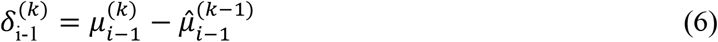

### FMRI Data Processing

FMRI data were collected on a Philips Achieva 3.0 T X-series scanner using a 32-channel head coil. For functional imaging, we used a T2*-weighted echoplanar imaging (EPI) sequence (voxel size, 2×2×3 mm^3^; field of view, 224×224 mm^2^; interslice gap, 0.6 mm; TR= 2800 ms; TE = 36 ms) to acquire 36 transaxial slices covering the whole brain. Pre- and post-processing of the fMRI data was done using the Statistical Parametric Mapping software SPM12 (Wellcome Department of Imaging Neuroscience, University College London, London, UK; http://www.fil.ion.ucl.ac.uk/spm) implemented in MATLAB R2022a (MathWorks). All fMRI images were first applied to slice time correction, spatial realignment, and normalization to the MNI template using the unified segmentation approach (Ashburner & Friston, 2005). Finally, normalized images were spatially smoothed using a Gaussian filter with a full-width half-maximum kernel of 6 mm. Data were high pass filtered at 1/128 Hz. For each participant, we conducted a first level general linear model (GLM). Events were time-locked to the onset of the presentation of the cue stimulus using stick functions and split into two regressors, one for *Staying* (no strategy switches) and the other one for *Switching* trials. For each of these two regressors, two parametric modulators of prior belief were defined. The first parametric modulator was prior belief about the sample-target contingency 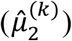. The second modulator was prior belief about the volatility 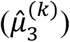, orthogonalized with respect to 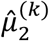. Please note that the absolute value of prior belief 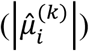 was used to enter the GLM as parametric modulators. Invalid trials (i.e., missing or late responses) were modeled separately. Furthermore, six head motion parameters, as estimated during the realignment procedure, were added as regressors of no interest to minimize false-positive activations due to task-correlated motion.

Using the GLM, we investigated prefrontal and thalamic responses when switching the decision strategy using the contrast ‘*Switching*> *Staying*’. The detailed data analysis for this comparison and corresponding results have been reported in our recent paper (Hummos et al., 2022). In current study, we primarily analyzed the main effect of parametric modulation by prior belief (both 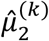 and 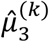) for all trials using the GLM. The respective t-contrast images of the modulatory effect for each subject were applied to the group-level one-sample *t* test (p < 0.05, family-wise error (FWE) corrected for the whole brain).

### Nonlinear Dynamic Causal Models (DCM)

In our previous study (Hummos et al., 2022), the fMRI GLM analyses revealed two prefrontal regions, i.e., right dlPFC and lOFC, together with the MD thalamus, which were all significantly modulated by decision switches (Fig. 1A). The Bayesian parameter averaging (BPA) across participants revealed that connections from lOFC to MD, from dlPFC to lOFC, as well as between dlPFC and MD in both directions were all significantly strengthened by strategy switches (Fig. 1B). In the current study, we show that the vmPFC reflects the value of the prediction, i.e., the prior belief about the sample-target contingency (see results section). Based on these findings, we next questioned how increased BOLD responses in the vmPFC, which were positively related to stronger prior belief, modulated the connection strength of the prefrontal-MD network, thus enhancing its computational efficiency and facilitating strategy switches. To test this, we constructed a nonlinear DCM including right dlPFC, right lOFC, right MD and vmPFC, and compared several alternative vmPFC modulations. For each brain region, subject-specific time series were extracted from the nearest local maximum within a sphere with a radius of 8 mm centered on each node’s group maximum. The first Eigenvariate was extracted across all voxels surviving *p* = 0.05, uncorrected, within a 4 mm sphere centered on the individual peak voxel. The resulting BOLD time series were adjusted for effects of no interest (e.g., invalid trials, and movement parameters).

The basic architecture of the model, shown in Fig. 1B and in Hummos et al., 2022, included the driving sensory (i.e., tactile) input directed to the dlPFC, as well as the four connections that were all significantly strengthened by strategy switches (i.e., dlPFC to lOFC, lOFC to MD, as well as between dlPFC and MD in both directions). With the nonlinear DCM we extended this model by the modulation of these connections through the activity in the vmPFC, which was driven by the trial-by-trial prior belief derived from the HGF model (Fig. 1D). We specified the model space with the modulatory influence of the vmPFC on different connections. More specifically, we tested the modulatory influence of prior belief related vmPFC on two out of the four significant connections within the dlPFC-lOFC-MD network, resulting in six models that were compared to each other (Supplementary Fig. S1A). The fixed-effects Bayesian model selection (BMS) was used to assess the most likely model among the six competing models. Parameters of the winning model were then summarized by Bayesian parameter averaging (BPA), which computes a joint posterior density for the entire group by combining the individual posterior densities. A posterior probability criterion of 90% was considered to reflect significant modulatory effect on the connections.

## Data and code availability

The data and code that were applied to assess the findings of this study are available from the corresponding author upon reasonable request.

## Acknowledgements

This work was supported by the Deutsche Forschungsgemeinschaft (DFG, German Research Foundation): Project number 122679504 -SFB 874 ‘Integration and Representation of Sensory Processes’ (to B.A.W. and B.P.), the DFG grant with the project number PL602/6-1’ Prefrontal-thalamic control of cognitive flexibility – from mice to humans’ (to B.P.) and National Natural Science Foundation of China: Project number 32200867 (to B.A.W.).

