## Supplementary figures and images for "Modulation of prefrontal couplings by prior belief-related responses in ventromedial prefrontal cortex"

### Supplemental Figure

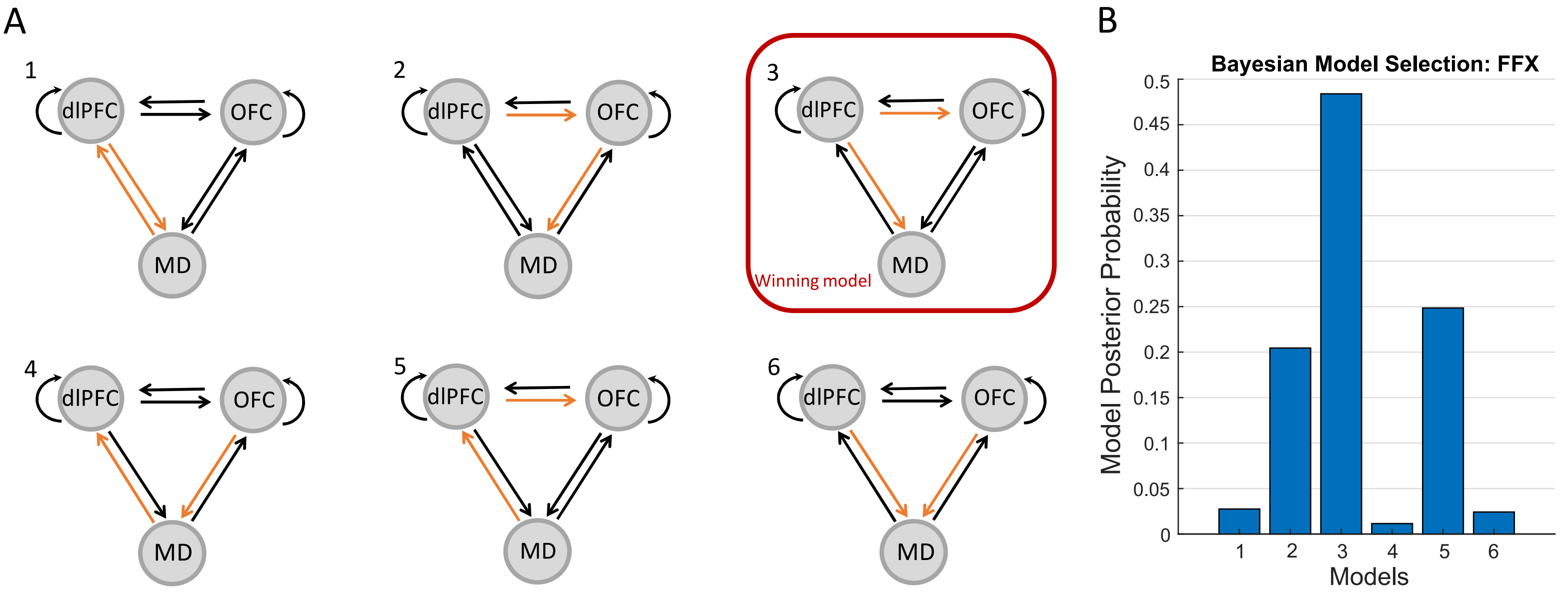
